# Two conceptions of evolutionary games: reductive vs effective

**DOI:** 10.1101/231993

**Authors:** Artem Kaznatcheev

## Abstract

Evolutionary game theory (EGT) was born from economic game theory through a series of analogies. Given this heuristic genealogy, a number of central objects of the theory (like strategies, players, and games) have not been carefully defined or interpreted. A specific interpretation of these terms becomes important as EGT sees more applications to understanding experiments in microscopic systems typical of oncology and microbiology. In this essay, I provide two interpretations of the central objects of games theory: one that leads to reductive games and the other to effective games. These interpretation are based on the difference between views of fitness as a property of individuals versus fitness as a summary statistic of (sub)populations. Reductive games are typical of theoretical work like agent-based models. But effective games usually correspond more closely to experimental work. However, confusing reductive games for effective games or vice-versa can lead to divergent results, especially in spatially structured populations. As such, I propose that we treat this distinction carefully in future work at the interface of EGT and experiment.

---

In my views of game theory, I largely follow Rubinstein (2012): game theory is a set of fables. A collection of heuristic models that helps us structure how we make sense of and communicate about the world. Evolutionary game theory (EGT) was born of classic game theory through a series of analogies. Given this heuristic genealogy of the field, it is usually alright to not worry too much about what exactly terms like strategy, player, or game really mean or refer to. I am usually happy to leave these terms ambiguous so that they can motivate different readers to have different interpretations and subsequently push for different models of different experiments. I think it is essential for heuristic theories to foster this diverse creativity: anything goes.

However, not everyone would agree with Rubinstein and me; some people think that EGT isn’t “just” heuristics. EGT is also directly empirically useful for questions in both mathematical oncology and the evolution of microorganisms. Microscopic experimental systems in which EGT has been useful include: *Escherichia coli* (Kerr et al. 2002), yeast (Gore, Youk, and Van Oudenaarden 2009; MacLean and Gudelj 2006), bacterial symbionts of hydra (Li et al. 2015a), breast cancer (Marusyk et al. 2014), pancreatic cancer (Archetti, Ferraro, and Christofori 2015), and lung cancer (Kaznatcheev et al. 2017b). But when we actually start doing experiments like these, it is no longer acceptable to be willy-nilly with fundamental objects of the theory: strategies, players, and games. The biggest culprit is the player. In particular, I think that a lot of confusion stems from saying that “cells are players”.

In classical game theory, the concepts of player, strategy, and game are intertwined but relatively straightforward. Players use a rational decision process to select strategies which are then mapped by the rules of the game to payoffs – the utility given to the players. Or, as I say more awkwardly in the first column of table 1: utility is given to a player based on its strategy which results from a rational decision process carried out by the player. All of this is summarized as the game. But, how does this classic picture translate to evolutionary games?

**Table 1:**
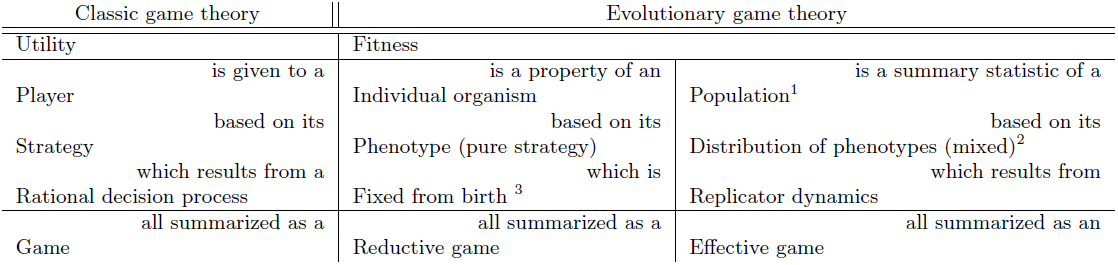
Defining evolutionary game theory by two different analogies to classic game theory.

| Classic game theory |  | Evolutionary game theory |  |
| --- | --- | --- | --- |
| Utility |  | Fitness |  |
|  | is given to a | is a property of an | is a summary statistic of a |
| Player |  | Individual organism | Population<sup>1</sup> |
|  | based on its | based on its | based on its |
| Strategy |  | Phenotype (pure strategy) | Distribution of phenotypes (mixed)<sup>2</sup> |
|  | which results from a | which is | which results from |
| Rational decision process |  | Fixed from birth<sup>3</sup> | Replicator dynamics |
|  | all summarized as a | all summarized as a | all summarized as an |
| Game |  | Reductive game | Effective game |

The easiest place to start is utility: this is almost always interpreted as changes in fitness. We can take this much as uncontroversial. But fitness itself has a complicated ontology and is part of a more general discussion in biological theory that reaches beyond EGT. Two competing interpretations are fitness as a property of an individual organism (reductive fitness leading to reductive games; see section 1) versus fitness as a summary statistic (or emergent property) of a population of subpopulation (effective fitness leading to effective games; see section 2). These two readings are summarized in columns two and three of table 1.

It is helpful to highlight the difference between these two interpretations with an analogy to physics. The setting of statistical mechanics mirrors the fitness for individuals view and defines properties like kinetic energy for individual molecules. Thermodynamics mirrors the effective fitness view and defines properties like temperate for ensembles of molecules. It simply doesn’t make sense to talk of the temperature of an individual molecule. Of course, in simple models like the ideal gas, temperature is just mean kinetic energy; this would correspond to an unstructured (inviscid) population in biology where the effective fitness of a (sub)population is just the average reductive fitness of the individuals that make up that subpopulation. But this ideal case seldom happens in nature. In general, there are many ways like recombination systems, spatial subdivision, and admixture in which structured populations depart from panmixis (mean-field). This can be very relevant to how we interpret games, as I discuss in section 3 in the context of spatial structure.

## 1 Fitness for individuals & reductive games

Since many evolutionary game theorists are computational modelers or think in terms of simulations and agent-based models, we take fitness as a property of an individual organism. In that case, we can define players as the organisms that receive the payoff from local interactions that happen between pairs of organisms (or more for multi-player games). The summary of this local interaction is what I would call the reductive game.

In a the most common EGT setting, what the organisms do in the game is fixed by their genes. Under this reductive interpretation players don’t alter their strategies. This makes it easy to present classic vs evolutionary game theory as two extremes on the spectrum of decision making. In classic game theory, players are unbounded rational decisionmakers. In evolutionary game theory, players are the most bounded possible: they make no decisions at all; their behavior is genetically fixed. Or, as I say more awkwardly in the second column of table 1: fitness is a property of an individual organism based in its phenotype, which is fixed from birth. All of this is summarized as the reductive game.

The proportion of agents in the population is then updated according to an evolutionary process like replicator dynamics. In particular, in the section I’ll discuss how two different implementations of reductive fitness can both give us replicator dynamics. In section 1.1, I will focus on fixed population sizes with fitness as probability to reproduce. In section 1.2, I will consider exponentially growing populations with fitness as number of offspring.

### 1.1 Moran: fitness as probability to reproduce

In a Moran process (Moran 1958; Taylor et al. 2004), we imagine that a population is made up of a fixed number N of individuals. An agent is selected to reproduce in proportion to their game payoff, and their offspring replaces another agent in the population, chosen uniformly at random. This gives us a very clear individual account of fitness as a measure of the probability to place a replicate into the population.

Traulsen, Claussen, and Hauert (2006) wrote down the Fokker-Planck equation for the above Moran process, and then use Ito-calculus to derive a Langevin equation for the evolution of the proportions of each strategy *x_k_*. The fluctuations in this stochastic equation scale with 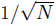 and so vanish in the limit of large *N*. This reduces them to a deterministic limit of the replicator equation in Maynard Smith form (Maynard Smith 1982), with the fitness functions as the payoff functions:

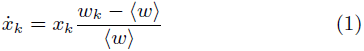

where 〈*w*〉 is the average fitness and the extra condition of 〈*w*〉 > 0 is introduced.^4^

Alternatively, we might be interested in directly getting the Taylor form (Taylor and Jonker 1978) of replicator dynamics:

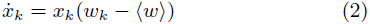

Traulsen, Claussen, and Hauert (2006) show how to achieve this, too. Instead of birth-death, they consider an imitation process. Two agents are selected uniformly at random, and individual if the payoff of the first individual is *w*_1_ and the second is *w*_2_ then the first copies the second with probability 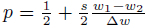 where Δ*w* is the maximum possible gap in the payoff of two agents in the model. With this version, they get the Taylor form, with the fitness as payoff, again. They are not the first to derive the Taylor form replicator equation from imitation processes. In fact, Schlag (1998) went further by showing that with the proportional imitation rule (only copy those that have higher payoffs, in proportion to how much higher the payoff is), you not only get the Taylor form replicator equation in a large population limit but also that this local update rule is optimal from the individual agent’s perspective in certain social learning settings.

### 1.2 Exponential: fitness as number of offspring

One of the biggest difference between ecological modeling in micro- vs. macro-organisms is that macro-organisms seldom have the opportunity to undergo exponential growth; they are almost always at carrying capacity. So what if we want to model this ecological difference – the fact that the total population grows or that the cell density in the Petri dish changes?

Populations don’t have to be constant to achieve replicator dynamics. Consider m types of cells with *N*_1_,…, *N_m_* individuals each, leaving *w*_1_,…, *w_m_* offspring each for the next generation. These offspring numbers *w_k_* could be functions of various other parameters. The population dynamics are then described by the set of m differential equations: 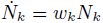 for 1 ≤ *k* ≤ *m*. Now, with *N* = *N_1_* +…+ *N_m_*, we can look at the dynamics of *x_k_* = *N_k_/N*:

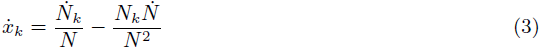

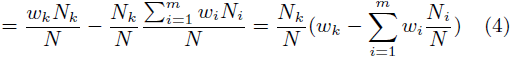

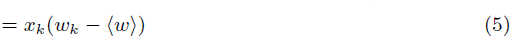

which is just the replicator dynamics. if the *w_k_* are functions of proportions then replicator dynamics can perfectly describe an exponentially growing population.

## 2 Fitness for populations & effective games

But fitness doesn’t have to be defined individually. An alternative perspective is to see fitness as defined only as a summary statistic or emergent property of populations. This is the perspective that makes the most sense when operationalizing fitness in microscopic systems; especially when using typical fitness measures like growth-rates. in that case, the player is the (sub)population that receives the payoff of fitness. The game then becomes the macroscopic coupling between (sub)populations made up of microscopic agents. it is even misleading to call this coupling an “interaction” since that suggest something too active and direct; as i show in section 2.5, the coupling could be as indirect as two populations feeding on a single resource in batch culture. This population-level description is what i call an effective game. Given its roots in operationalization of microscopic systems, the effective games can be measured directly and we recently developed a game assay for this purpose (Kaznatcheev et al. 2017b).

This perspective has some curious consequences. Since the players are populations, the individuals organisms – or behaviorally identical classes of them – are the strategies. The distribution of phenotypes in the population is then interpreted as a mixed strategy. The player is not static but carries out a ‘decision process’ specified by the rules of the evolutionary dynamics. This is usually described by the replicator equation. Make what you will of the correspondence between replicator dynamics and Bayesian inference, reinforcement learning and other forms of rational decision making (Arora, Hazan, and Kale 2012; Borgers and Sarin 1997). To me, this seems like both a closer correspondence to the aspirations of classical game theory and easier to link to experiment.

To summarize as in the third column of table 1: fitness is a summary statistic of a (sub)population based in its distribu-tion of phenotypes, which is updated according to replicator dynamics. All of this is summarized as the effective game.

What does this mean for “cells are players”? For the reductive game, each individual cell is a player. For the effective game, the population of cells is a player.

### 2.1 Replating: fitness as fold change

Consider the following idealized protocol that is loosely inspired by Archetti, Ferraro, and Christofori (2015) and the *E. coli* Long-term evolution experiment (Lenski et al. 1991; Ribeck and Lenski 2015; Wiser, Ribeck, and Lenski 2013). We will (E1) take a new petri dish or plate; (E2) fill it with a fixed mix of nutritional medium like fetal bovine serum; (E3) put a known number 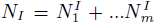 of *m* different cell on the medium (on the first plate we will also know the proportion of A and B in the mixture); (E4) let them grow for a fixed amount of time Δ*t* which will be on the order of a couple of cell cycles (keeping us in the growth phase); (E5) scrape the cells off the medium and measure the final numbers 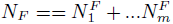; and (E6) return to step (E1) while selecting *N* cells at random from the ones we got in step (E5) to seed step (E3).

From comparing steps E3 and E5, we can get the experimental population growth rates as:

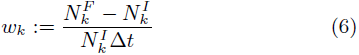

this can be rotated into a mapping *N_I_* ↦ *N_F_* given by 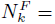 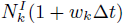.

From defining the initial and final population sizes 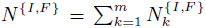, we can compare the initial and final proportions of each cell type 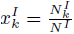 and

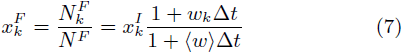

where 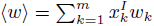.

So far we were looking at a discrete process. But we can approximate it with a continuous one. In that case, we can define 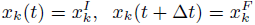 and look at the limit as Δ*t* gets very small:

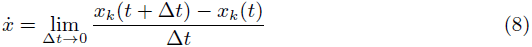

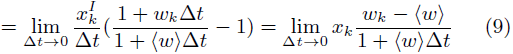

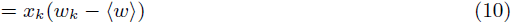

We recover replicator dynamics as an explicit experimental interpretation for all of our theoretical terms.

We didn’t make any assumptions about if things are inviscid or spatial; if we are talking about individual or inclusive fitness; or, if we have growing populations in log phase or static populations with replacement. All of these microdynamical details that we spent so much time debating about are simply buried in the definition of experimental fitness. More importantly, we provided a precise description of how we will measure this quantity. This allows us to hide the pedantics of microdynamics inside of how we measure. Somehow, this might seem unsatisfying in that we just named something but aren’t testing anything. This is where we can begin to twist our experimental knobs a bit to start understanding our system, as I do in section 2.3.

If we are able to peek inside the system more, as – for example, Kaznatcheev et al. (2017b) do with time-lapsed mi-croscopy – then we can also replace fold-change by more specific measurements like inferred growth rates. An advantage is that the goodness-of-fit of exponential models can provide a good estimate of the error associated with these measurements. But the cost is a slightly more specific set of assumptions on the microdynamics of our system. However, since many models can lead to transient exponential growth curves for various microdynamic implementations, these assumptions still don’t have to be as stringent as the definitions in section l.2.

Does this mean that just by playing with definitions and writing down the replicator equation I have shown that any replating experiment follows replicator dynamics? That would be ridiculous. But we aren’t far. To actually establish this, we need only one more piece: we would need to show that the the replatings have the Markov property. In other words, that if the protocol is kept fixed then from one plate to another then the proportion of cells (and maybe a little noise) is all we need to specify the proportion of cells that we will have at the end of the grow phase; there is no dependence of plates that are further back in time. Of course, there are many ways this Markov property might break down.

The easiest way to break the Markov property is if the state-space we consider is insufficient. For example, we assume that the ‘pure’ cultures we start with all have the same strategy. What if the culture is heterogeneous in terms of their game behavior? In that case, for a two-strategy game, we need to treat the game as not between type A and type B but between types A, A’ and B where we don’t have an experimental procedure to differentiate between type A and A’. Alternatively, the extra variable we need to add to the state space might not be an extra cell (sub)type but something like a toxin that survives between replatings.^5^

Of course, expanding the state space might be futile in some cases. The classic examples of this would be cellular or epigenetic memory, certain kinds of phenotype switching, or rapid mutations. In that case, our state space will expand with each replating and it is better to abandon replicator dynamics and adopt some technique that was built for handling historicity. To find this out, though, we would need to first design careful experimental protocols and run our cells and see the long-term behavior breaking the Markov property. Alternatively, if we are lucky and nature cooperates by granting our protocol the Markov property then we will find ourselves in the fortunate position of having a relatively simple operationalist theory of evolutionary games.

### 2.2 Measuring the gain function directly

We don’t necessarily need to measure separate fitness functions for each cell type. It is more important to know the fitness differences, which we can measure directly instead.

Suppose the proportion of cell line A in a mixture is *p*, with cell line B making up 1 – *p*, and the fitnesses of the cells are *W_A_* (*p*) and *W_B_* (*p*), respectively. Then the replicator dynamics are given by:

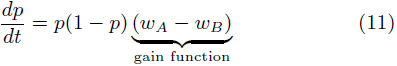

Our goal in identifying the game is to measure this gain function (see Peña, Lehmann, and Nöldeke (2014) and Kaznatcheev et al. (2017a) for their importance): the increase in growth rate from ‘switching’ from strategy A to B with *p* held constant. For this we will use a simple calculus trick: consider the log-odds 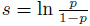. Then

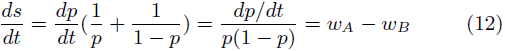

By looking at the log-odds of p instead of just *p*, we have ‘factored out’ the logistic growth part of the equation. Now, to measure the gain function, we just have to measure the derivative of *s*. Unfortunately, experimentalists do not have a derivative detector in the lab, so we have to approximate the derivative by looking at the change in *s* over a short period of time.

In the case of the basic replating experiments you have a natural discretization of time: *p*_in_ can be the proportion of type-A cells at the start of your experiment, and *p*_out_ can be the proportion of type-A cells when you replate. You can then run the experiments for several initial values *p*_in_ and plot your results as 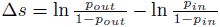 versus *p*_in_. This graph is your gain function.

Here we run into the important question of “how short is short enough?” If we run the experiment for too short of a time then the change in *p* will be overwhelmed by the measurement noise, but if we run for too long before measuring *p*_out_ then it doesn’t make sense to say we are measuring the derivative. To overcome this, we have to focus on the tangent line of our function as a local linearization. If we have access to high-resolution in time data of the growth during our preliminary experiments then we can plot the resulting growth curve as we would normally: proportion versus time. We then want to take the biggest time-window on which this growth curve is well-approximated by a constant-fitness logistic growth. This logistic growth is the linearization of our function, and as long as it is in approximate agreement, we can say that we are measuring the derivative.

Since we are considering experimental data, it is important to look at the errors associated with our measurements. I don’t mean the variance between different runs in different Petri dishes, although that is also important, but the accuracy of the proportions of our initial seeds and the precision of our measurements. There are several ways we could propagate the errors from *p* to *s*, but as an estimates:

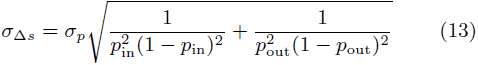

So the error is amplified by 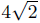 near *p*_in_ = *p*_out_ = 0.5 and the amplification increases as the proportions approach 0 or 1. For example, for *p*_in_ = *p*_out_ = 1/*m* and large *m*, it becomes approximate a factor of 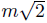. Of course, each experimental set up will serve as a different meter stick, and we will need to do the metrology more carefully for each.

### 2.3 Partitioning fitness with mixed media

For a conditioned medium experiment, a petri-dish is filled with a liquid containing some food (the medium) and then cells of type X from some established culture are introduced onto this medium. They spend some fixed amount of time in that dish, consuming a portion of whatever is in the medium and potentially releasing their own chemicals into the liquid. After a relatively short amount of time – something on the order of the cell cycle – the cells are separated from the medium, usually by running the mixture through a filter that is fine enough to catch cells but let macro-molecules through. This gives us the post-X medium in a flask, which we pour out into a fresh petri-dish and now introduce a known number of cells of type Y onto it. Again, we grow this mixture for a fixed amount of time on the order of a cell cycle and then the cells. This allows us to work with monotypic cultures, which can be helpful if the tools for marking or sorting cells by type are unavailable.

We can extend the conditioned media experiments to mixed media experiments: take a beaker of conditioned medium from cells of type A and another from cells of type B and mix them in some proportion *p*. Now we can grow cells of type X on this mixed medium and the ratio of cells that we introduced to the cells we count at the end will give us the fitness of X as a function of the proportion *p* of type A medium in the A-B mix. By considering different values of mixed media *p* added to monoculture, we can do a mixed-media version of the game assays described in section 2.1 and Kaznatcheev et al. (2017b).

This gives another operationalization of fitness: experimental medium-mediated fitness. When we grow cells X on media conditioned by a proportion *p* of type A conditioned media in the A-B mix then we can call the corresponding ratio between initial and final cells as 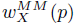.

Now, if we compare the experimental medium-mediated fitness to the experimental fitness *w_x_* (*p*) that we defined above to 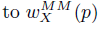 then for a short enough replating time, we would expect the two numbers to be nearly identical if the interaction between cells is primarily mediated by the medium: 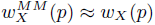.^6^

In general these two functions might be very different. In that case we will have made more than a measurement. We will have discovered that the experimental fitness function partitions into a direct interaction part – that could come from direct predation, very short-term or short-range media-based signals, spatial competition, or many other microdynamical effects – and a medium-mediated part: *w_x_ (p)* = 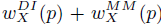.^7^ In other words, we started by trying to eliminate all reductionist basis from our work, and yet learned something about the microdynamical properties of our system. Although coming at it from the operationalist perspective forces us to be much more humble and reserved about our conclusions. If we come up with other experimental knobs to fiddle with then we can find further partitions and thus more nuance for the operationalist meaning of the fitness function.

### 2.4 Choosing units of size for populations

So far, we mostly considered populations as numbers of individuals, organisms, or cells. But what is so special about the number of cells? In this subsection, I want to question the reasons to focus on individual cells (at the expense of other choices) as basic atoms.

So, let’s look at what we could mean by ‘size of population’. The obvious definition is number of cells, and if all we did was *in silico* simulations then it is the definition we could stuck to. Especially for agent-based models, it is very tempting to have cells as your agents and building everything up around them. But consider two populations that have the same number of cells and everything else is equal, *except*…

1. *…the cells in the first population are metabolically twice as active as cells in the second population.* In this case, the more active cells can easily strain their environment more, as they use more resources to fuel themselves. If your limiting resource in the petri dish is growth medium then the more metabolically active cells will consume more of it.^8^ With slower metabolic activity, the cell becomes less of an effect not only on its own future, but also on other cells it interacts with – for example, by moving around less or releasing fewer cytokines and thus interacting with fewer other cells. In this case, the more natural set of physical units might be the power consumption in terms of watts-used or ATP-used. This might be more compatible with the metabolic theory of ecology.(Brown et al. 2004)
2. *…the cells in the first population are twice as big as cells in the second population*.^9^ From the point of view of games mediated by things like diffusive factors or cellcell contact, the bigger cells will have more area to absorb/release factors or to contact other cells. In we are working *in vitro*, larger cells also exhaust the limiting factor of free space quicker than small cells.^10^ On top of this, size feeds back into the first point, with larger cells usually doing more things metabolically and in terms of other activity. In this case, the more natural set of physical units might be area-covered.^11^

Without individual cells to ground us, reductionist story telling becomes more difficult; something that can be both a plus or a minus:

On the one hand, it is hard to imagine how 10 watts-used by cancer interacts with 10 watts-used by fibroblasts, instead we are forced to make these measurements experimentally. Since these measurements are almost always at the level of populations, we don’t feel a need to make sense of them in reductionist terms of how a single watt-used interacts with another watt-used. You might have noticed that even the word ‘interaction’ felt awkward in the last two sentences. Watt or area use invite us to recognize the importance of both the size of the other population and the environment more generally. This makes it easier to notice evolutionary games with only indirect interactions.

On the other hand, these more abstract units and operationalist perspective can hinder the imagination, and it can often become more difficult to explain the work or to design new experiments. The alternative units I considered here also obscure the discrete nature of cells, a discreteness that is essential to life and can have significant side-effects on model conclusions (for discussion, see Durrett and Levin (1994) and Shnerb et al. (2000)). Although in the case of replicator dynamics, it is not clear how much this would matter since they already ignore the discrete nature of life through the use of ODEs.

### 2.5 Effective games without interactions

Let *p* be the proportion of A-cells in the population, and *y* be the nutritional content of the medium – normalized so that the most nutrient rich mix possible has *y* = 1 and distilled water has *y* = 0. For each cell we will have some (analytic) feeding function *f_A_* (*y*) and *f_B_* (*y*) which translates between the nutritional content of the medium and the organism’s fitness, such that:

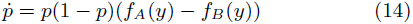

It is important to note that *f_A_* and *f_B_* are functions of y and completely independent from *p*. At this point, we might be tempted to stop by saying that since experimental step (E2) uses a fixed mix of nutritional medium, we can just treat *f_A_* (*y*) – *f_B_* (*y*) as a constant and thus (excluding the neutral case) we will always have the population converge to all-A or all-B depending on the sign. We will see no (non-trivial) evolutionary game dynamics. This is the standard intuition behind Gaus’ exclusion principle: two species cannot co-exist on a single abiotic resource (Hardin 1960).

But stopping here would be a bit disingenuous. The reason that we have to renew the medium on each cycle of the experiment is because it gets consumed between the replating. Further, the rate of consumption might differ between the two cell types. suppose that each cell type consume the nutrients at rate 2*k_A_* and 2*k_B_*, such that if *y*_in_ was our initial level of nutrients in step (E2) then our final level it step (E5) is *y*_out_ = *y*_in_(1 - 2*k_A_x* – 2*k_B_* (1 – *p*)).

We can assume that the cell cycle is significantly slower than the metabolic cycle, and start working with our average consumption (calling *y*_in_ just *y*): 〈*y*〉_*p*_ = *y*(1-*k_B_* +*p*(*k_B_*-*k_A_*)).

Now, our dynamics become:

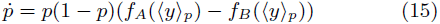

and suddenly our gain function is no longer independent of *p*.^12^ That is the main slight-of-hand, but let’s take the trick to its conclusion.

We will expand the gain function, noting that we picked analytic feeding functions, so 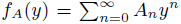 for some sequence {*A_n_*} and similar for *f_B_*(*y*) but with {*B_n_*}:

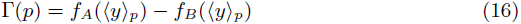

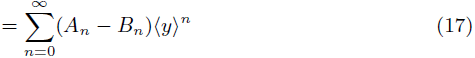

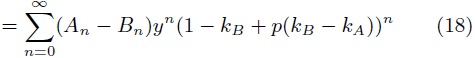

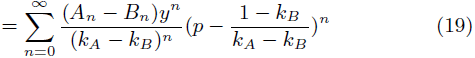

The last line is the power series of some analytic function Γ(*p*) with coefficients 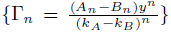 around the point 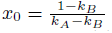. In particular, given any desired (analytic) gain function Γ(*p*), there is some choice of feeding functions *f_A_* and *f_B_* (thus, their corresponding coefficients {*A_n_*} and {*B_n_*}) and *k_A_* and *k_B_* such that the population follows identical dynamics. In other words, in this experimental set up, we can recreate any evolutionary game dynamics without having the cells interacting but just based on how they turn nutrition into reproduction. in particular, we can implement games like Hawk-Dove to have co-existence of A and B on a single abiotic resource, thus violating the competitive exclusion principle.

## 3 Spatially structured populations

Spatial structure in game theory is so important that a particular approach to it even has its own sub-field name: evolutionary graph theory(Lieberman, Hauert, and Nowak 2005; Maciejewski and Puleo 2014; Shakarian, Roos, and Johnson 2012; Szabό and Fath 2007). Durrett and Levin (1994) and Shnerb et al. (2000) provide a particularly good demonstration of how much spatial structure and stochasticity can matter as they build from mean-field approaches (of which the inviscid replicator dynamics with which we started is an example) to patch models of discrete individuals to reaction-diffusion equations to full-fledged interacting particle systems. We know that space can promote cooperation (Nowak and May 1992; Ohtsuki et al. 2006), or inhibit it (Hauert and Doebeli 2004), or complicate the whole discussion around it (Killingback and Doebeli 1996). Spatial structure can so drastically change the nature of the reductive game such that the mean-field analysis in completely inapplicable. in section 3.1, i will consider how we might work from the bottom up, by transforming a reductive game through an incorporation of spatial structure. But in section 3.2, I’ll consider the reverse direction: extracting the contribution of space from an effective game that would normally swallow-up or abstract the spatial structure into measurement.

### 3.1 Approximating spatial structure

A typical study of space in evolutionary game theory will start with the reductive game and then simulate that interaction over a model of space to show a surprising difference in dynamics between the spatial model and the mean-field. Exceptional works like that of Ohtsuki and Nowak (2006) (more recently, Nanda and Durrett (2017)), provide a general method for combining a reductive game with spatial structure. Although in its original presentation, Ohtsuki and Nowak (2006) focused on dynamics on *k*-regular random graphs, i think it can be useful to frame their work as a general first-order approximation of an arbitrary spatial structure.

First, let me present replicator equation as a zeroth-order approximation of spatial structure. Without knowing anything about spatial structure, the roughest guess we can make of the probability of interacting with another agent is just to say that we sample agents from a distribution given by their proportion in the population – a mean-field approximation. This gives us a utility function for agents of type *i* as [*G_x_*]*_i_* where *G_ij_* is the payoff of an agent of type *i* interacting with an agent of type *j* and *x* is a vector of proportions of agents of each type. From here on in, our hands are tied and the math forces us to replicator dynamics: 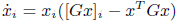.

But the perfect sampling used in the calculation of fitness effectively makes interactions global. To overcome this, we can say that instead of the fitness being the mean-field [*G_x_*]*_i_*, we instead sample *M* interaction partners from the distribution given by *x* and use these local interaction groups for our fitness calculation. This would be our 0.5th-order approximation. in this case, Hilbe (2011) showed that the result is still replicator dynamics (although with a different time-scale that is irrelevant to the functional form) but with a modified payoff matrix 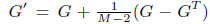. Where *G^T^* is the matrix transpose of *G*. This is a great way to reintroduce some local effects, but the groups of *M* agents are constantly re-sampled and fitness-competition still happens at a global level; in other words, there is no spatial structure. Hence the 0.5 and not 1.

To get things completely localized, we will assume a fixed population size, and make our replication procedure more explicit to make a first-order approximation of spatial structure. Since the population size is fixed, we can only get a new agent if an old one dies, this gives us a great way to localize. Once a focal agent dies, there is some neighborhood of the focal agent with *k* agents that compete for the focal spot. Now, we can have some extra information, instead of just keeping track of the proportion of agents of type *i* given by *x_i_*, we can keep track of neighbors. More explicitly, we will use pair-approximation (Matsuda et al. 1987; Van Baalen 2000) to keep track of proportion *x_ij_*: the probability of seeing an agent of type *i* in the neighborhood of an agent of type *j*.

This tells us who is competing for for the vacated spot, but it doesn’t let us calculate their fitness because we would need to know the probability *x_ijk_* of seeing an agent of type *i* near an agent of type *j* and *k*. To update that probability, we would need to know more long range effects like *x_ijkl_*, etc. Hence, for a first-order approximation, we truncate the series here and approximate the further effects by saying that *x_ijk_* = *x_ij_*.^13^ Working under these assumptions, Ohtsuki and Nowak (2006) showed that we still get replicator dynamics but with a modified payoff matrix 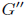 given by:

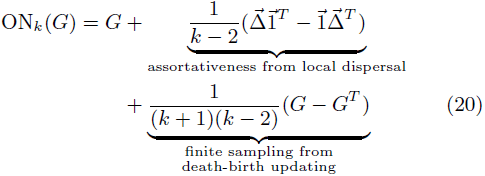

where vector Δ which is the diagonal of the game matrix *G*, i.e. Δ_*i*_ = *G_ii_* and 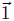 is the all ones vector; thus 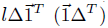 is a matrix with diagonal elements repeated to fill each row (column). Note that we have *k* agents competing for a spot, and each one samples *k* – 1 other agents (since one spot they are neighboring was just vacated) so *M* = *k*(*k*-1). Thus, the last term is Hilbe (2011) finite sampling effect and the first term is the spatial structure.

To restate: the effective game 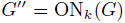 then has the same mean-field replicator dynamics as the reductive game *G* carried out on the spatial structure. This allows us to use our tools of evolutionary game theory to analyze the transformed game and thus learn something about the system implemented by the reductive game and spatial structure. We can think of this approach as bottom up: start with the reductive game, find the corresponding effective game (or other description of macroscopic population-level dynamics) and then use these as a prediction to compare against observed phenomena.

As an example of the typical study of space in EGT, consider Kaznatcheev, Scott, and Basanta (2015). Here, we started with the reductive Go-vs-Grow game between invasive (INV; Go) and autonomously growing (AG; Grow) cells (Basanta, Hatzikirou, and Deutsch 2008). We wanted to know how the game was effected by different spatial structure in the bulk versus a static boundry of a tumour, so we transformed it according to the Ohtsuki and Nowak (2006) transform:

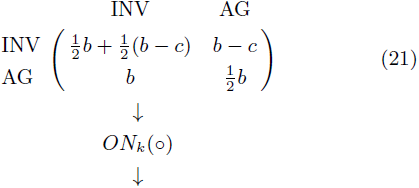

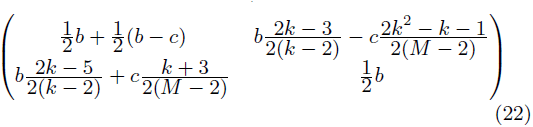

where *M* = *k*(*k* – 1).

Unlike the reductive game in equation 21, the effective game in equation 22 allows for a fully invasive tumour when 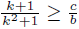. From this, we drew the surprising conclusion that a tumour can have much more invasive phenotypes at the boundary than the bulk.

### 3.2 Operationalizing spatial structure

But when we apply the typical pipeline: how do we know that the local interactions of the reductive game are the right ones to start with? In the example above, how do we know that the reductive Go-vs-Grow game is given by equation 21? For macroscopic systems like human or other large animals, we might be able to directly observe or maybe even design the reductive game. In microscopic systems like cancer, however, we tend to guess these games from intuitions acquired by looking at population level experiments. Unfortunately, these experiments seldom explicitly account for the effect of their spatial structure. Hence, they are actually intuitions about the effective game that we then feed into our models as the reductive game. This is the common confusion about spatial structure in microscopic systems. We are taking a game from a top-level view, feeding it into the bottom level, getting a different result at the top-level and then publishing that surprising conclusion.

This is backwards. At best, it is just telling us that our intuitions about the game were wrong – since correct intuitions about the reductive game should yield the observed effective games. At worst, this is a type-errors and thus incoherent: we are feeding in an effective game where we should be putting a reductive game.

Instead, I propose starting with our measurement of effective games and pushing it down with an operationalization of spatial structure. We can achieve this, by giving an experimental definition of the local environment of cells. This local perspective might be very different from the perspective that an experimenter has of the system as a whole.

Let’s suppose we are considering a system with two possible strategies A and B then we can define the following functions:

- Let 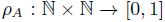 be the distribution over number of type A and type B partners that a player of strategy A encounters; i.e. *ρ*(*k_A_, k_B_*) is the probability of encountering *k_A_* many players of type A and *k_B_* many players of type B during the timescale relevant to the calculation of local fitness. This is similar to *p_ij_* in our discussion of the Ohtsuki-Nowak transform.
- Let 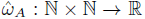 be the local fitness function for an agent of type A; i.e. *ŵ_A_(k_A_,k_B_*) is the local fitness of an agent of type A that encountered *k_A_* many players of type A and *k_B_* many players of type B during the timescale relevant to the calculation of local fitness.

Define *ρ_B_*, *ŵ_B_* analogously for strategy B. This allows us to write down the replicator dynamics at the level of the whole population as 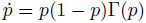 where the gain function Γ(*p*) is given by:

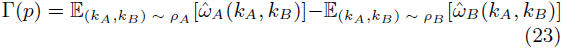

Since *ρ*{_*A*_,_*B*_} depends on *p*, let us name this mapping 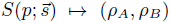 and introduce an extra state vector 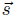 which might also change with time according to some general relationship: 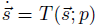. This relationship between *p*,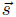 and *ρ* is meant to capture the functional role of space or any other discrepancy between the local and global perspectives. The hope is that in practical settings, 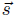 is simple or non-existent or the dynamics of 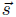 can be decoupled from the dynamics of *p* by something like a separation of timescales, like what happens from using weak selection in the derivation of Ohtsuki and Nowak (2006).

Now, suppose that we were able to find a system where the equations for 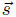 can be decoupled from *p*. In that case, this formalism is that we can start to operationalize space. The first step to measure the gain function Γ(*p*) as described in section 2.2. But measuring *S* is more difficult because it is encoding much more information about the microdynamical structure. A good first guess might be to take something like a structured core biopsy and define an interaction radius *r*. Then go through taking each cell as a focal agent and count how many cells of type A and B are within distance *r* of the focal agent. The result is an empirical estimate for *ρ_A_*, *ρ_B_*. Repeat for different initial *p* to get as many points of function *S* as desired. Clearly, if *r* is taken as the diameter of the slide then *S* will be basically an identity map since 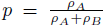 in that case. At the other extreme, taking *r* as less than a cell radius will make *S* into a constant map with *ρ_A_* = *ρ_B_* = 1. For intermittent values of *r*, we will potentially have a variance in different local densities for each focal agent, and picking a good *r* will depend on trade-offs between the level of error introduced by this variance versus the level of error that’s introduced in the propagation from raw gain function to game.

Unfortunately, just like with other effective concepts, this operationalized of spatial structure might not always have clear microdynamic interpretations. However, it does allow us to go one step closer to experimentally understanding the effects of spatial structure on populations without confusing effective and reductive games.

## Footnotes

1 Since we are primarily interested in fitness differences, it is usually subpopulations and not the population of all organisms.

2 If the fitness functions are linear then interpreting the distribution of phenotypes as a mixed strategy is straightforward. However, fitness functions are not necessarily linear; see Li et al. (2015b) for a clear example. In the case of a fitness function given by a *m*-order polynomial, we can still interpret the distribution of phenotypes as a mixed strategy in an (*m* + 1)-player game.

3 Fixed from birth is the typical case but it is also natural for the evolutionary game theorists to gradually build up the decision making capacities of the player (McNamara 2013). For example, tag-based models can be seen as taking a minimal step upwards by allowing the focal to decide to cooperate of defect conditioned on the tag of the alter. Although this characterization behavior is simple, it can allow for a rich analysis as a form of minimal cognition (Beer 2003) and we can associate a cognitive cost for this extra decision-making ability (Kaznatcheev 2010). Or push this genotype to behavior map even further (Kaznatcheev, Montrey, and Shultz 2014) by having evolving agents act rationally on their evolved perceptions of the game payoffs and (potentially-biased) estimates of other’s probability to cooperate (for this direction, see also Masel (2007)).

4 In the limit of weak-selection, this form is equivalent to the Taylor form. Even without weak-selection, the two systems of equations differ only by dynamic time rescaling and thus have the same fixed points, orbits, and paths. If we only care about this in our analysis then we can use the equations interchangeably.

5 This can be done by binding to the surface of the cells – although experimentalists often ‘wash’ cells to avoid this specific efiect – or by being absorbed without harm by one cell type to only be actively or passively pumped out of them later and affect the other cell type – a typical example in oncology might be a chemo-resistant cell pumping out the drug after replating and poisoning it’s non-chemo resistant partners.

6 Although even here, we might want to allow for some low complexity interpolation between the two functions, like proportionality, log-proportionality, or logit-proportionality and accept the approximate equality if the two functions are equal under this comparison instead of strict numerical equality which would correspond to the simplest map: identity.

7 Again, the best way to partition might be something other than ‘+’ (that corresponds to identity or proportionality) and we might use another low complexity function to partition like log (*w_x_ (p)* = 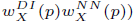 or logit 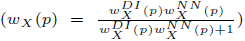. Whatever choice is made, should be consistent with the choice for approximation in ^6^ so that the domain to which the different ws map have a consistent mathematical structure.

8 The extreme case of this is cells that are completely inactive or maybe even dead. This doesn’t come up as much in simulation, since I can just cleanly remove dead agents.

9 In the case of cancer, the tumor corresponding to the more voluminous population would be much more burdensome to the patient. In fact, tumor burden is often measured and reported as volume in x-ray or other imaging. The number of cells in the tumor is then inferred from these volumetric measures by assuming (or measuring outside the body) the size of a typical cancer cell.

10 The importance of area has come up in thinking about prostate cancer metastases to the bone. (Araujo et al. 2014) Osteoclasts and osteoblasts, take up drastically different amounts of area on the bone, and they are only of significant consequence to the model if they are in contact with the bone (else they are not remodeling it). Area-On-Bone becomes the important variable here.

11 Of course, we could try to express the above in terms of individual cells by converting back and forth between numbers of cells and watts-used or area-covered. Practically, this would mean finding a conversion factor which amounts to a measure of how much power or area a typical cell uses. But in doing so, we have swept some amount of heterogeneity under the rug – after all, each cell takes up a different amount of space or uses a different amount of energy, especially when facing new circumstances like chemotherapy – and it is not clear what useful thing we got in return.

12 It might be helpful to switch to our prior notation by noticing that in this case *W*_{*a,b*}_(*p*) = *f*_{*A,B*}_(〈*y*〉_*p*_)

13 Since we assumed that the neighbors of the perished agent i are drawn from the same sort of distribution as the neighbors’ neighbors, we have ignored extra correlations that might arise from looking out to distance two or more, hence the first order nature of the approximation. Of course, like any first-order approximation, if higher-orders are important then it will not agree with experimental data. However, if we just look at data without a first-order theory then we wouldn’t even know that higher-order terms are important. Thus, the first-order approximation is always a good first step; if empirical results contradict it then at least we know where to look, second-order and higher correlations in the distributions of neighbors.

